# Necroptosis and apoptosis contribute to cisplatin and aminoglycoside ototoxicity

**DOI:** 10.1101/332031

**Authors:** Douglas Ruhl, Ting-Ting Du, Jeong-Hwan Choi, Sihan Li, Robert Reed, Michael Freeman, George Hashisaki, John R. Lukens, Jung-Bum Shin

## Abstract

Ototoxic side effects of cisplatin and aminoglycosides have been extensively studied, but no therapy is available to date. Sensory hair cells, upon exposure to cisplatin or aminoglycosides, undergo apoptotic and necrotic cell death. Blocking these cell death pathways has therapeutic potential in theory, but incomplete protection and lack of therapeutic targets in the case of necrosis, has hampered the development of clinically applicable drugs. Over the past decade, a novel form of necrosis, termed necroptosis, was established as an alternative cell death pathway. Necroptosis is distinguished from passive necrotic cell death, in that it follows a cellular program, involving the receptor-interacting protein kinases 1 and 3 (RIPK1 and 3). In this study, we used pharmacological and genetic intervention to test the relative contributions of necroptosis and caspase-8-mediated apoptosis towards cisplatin and aminoglycoside ototoxicity. We find that *ex vivo*, only apoptosis contributes to cisplatin and aminoglycoside ototoxicity, while *in vivo*, both necroptosis and apoptosis are involved. Inhibition of necroptosis and apoptosis using pharmacological compounds is thus a viable strategy to ameliorate aminoglycoside and cisplatin ototoxicity.

**Significance statement:** The clinical application of cisplatin and aminoglycosides is limited due to ototoxic side effects. Here, using pharmaceutical and genetic intervention, we present evidence that two types of programmed cell death, apoptosis and necroptosis, contribute to aminoglycoside and cisplatin ototoxicity. Key molecular factors mediating necroptosis are well characterized and druggable, presenting new avenues for pharmaceutical intervention.

## Introduction

The clinical application of the chemotherapeutic agent cisplatin, used in the treatment of various types of solid tumors (Einhorn, 2002; Burdett et al., 2015), is dose-limited due to nephrotoxicity and ototoxicity (Karasawa and Steyger, 2015). Cisplatin results in the death of sensory hair cells, but the underlying mechanisms are still poorly understood (Sheth et al., 2017). Evidence suggests that cisplatin exerts its cytotoxic effect in hair cells through binding to cellular DNA, causing transcriptional inhibition and cell cycle arrest (St. Germain et al., 2010), generation of reactive oxygen species, followed by mitochondrial damage and activation of apoptosis (Clerici et al., 1996; Kopke et al., 1997), involving the activation of MAP kinase pathways and caspases (St. Germain et al., 2010). The clinical use of aminoglycoside antibiotics is also hampered by its ototoxic side effects. Aminoglycosides enter hair cells through the mechanotransduction channel, accumulate in hair cells (Richardson et al., 1997; Marcotti et al., 2005) and cause oxidative stress (Priuska and Schacht, 1995), and initiate cell death through both caspase-dependent and independent mechanisms (Cunningham et al., 2004). Although certain pathways are specific for cisplatin or aminoglycoside ototoxicity, the two pathologies share remarkable commonalities in the stress response they elicit in sensory hair cells: For example, both elicit oxidative stress (Lautermann et al., 1995; Clerici et al., 1996; Hirose et al., 1997; Kopke et al., 1997; Dehne et al., 2000), activate p53 (Zhang et al., 2003; Coffin et al., 2013; Benkafadar et al., 2017) and the JNK pathway (Wang et al., 2004; Schacht et al., 2012; Francis et al., 2013). In addition, our lab previously demonstrated that both aminoglycoside- and cisplatin-induced ototoxicity is correlated with inhibition of cytosolic protein synthesis (Francis et al., 2013; Nicholas et al., 2017). Despite scores of compounds reported to protect against cisplatin and aminoglycoside-induced ototoxicity, it is a concerning reality that none of the described compounds have entered clinical trials yet. The stubborn intractability of cisplatin and aminoglycoside ototoxicity is likely caused by the fact that a multitude of stress pathways contribute to the pathology. Instead of attempting to block all upstream pathways involved, strategies to interfere with the final execution of cell death, by blocking programmed forms of cell death, could be a more tractable strategy to prevent ototoxicity. Caspase-mediated apoptosis is the most cited cell death pathway associated with ototoxicity, mainly affecting hair cells, stria vascularis and spiral ligament and spiral ganglion cells (Alam et al. 2000, Liang et al. 2005, Watanabe et al. 2003). Indeed, pan-caspase inhibitors such as zVAD-FMK effectively prevent ototoxin-mediated hair cell loss in explant studies (Forge and Li, 2000; Cheng et al., 2003). In contrast to the *ex vivo* studies, the mode of cell death *in vivo* is thought to involve both apoptotic and necrotic cell death (Forge and Li, 2000; Matsui et al., 2003; Okuda et al., 2005; Jiang et al., 2006). Necrosis in its classic form lacks any specific molecular mediators, hence is intractable for molecule-specific inhibition. Studies in the past 15 years, however, have uncovered an alternative, programmed form of necrotic cell death termed necroptosis that is accessible for pharmacological intervention using a class of small-molecule inhibitors called necrostatins. Together with various mouse models that enabled loss-of-function studies for mediators essential for necroptosis, including the receptor-interacting protein kinases RIPK1 and RIPK3 (Weinlich et al., 2017), this has allowed molecular dissection of the necroptosis pathways.

Here, we tested the involvement of caspase-8-mediated apoptosis and RIPK1/3-mediated necroptosis in kanamycin and cisplatin ototoxicity. Using mice with null mutations in Ripk3 and/or Caspase-8 (Salmena et al., 2003; Newton et al., 2004), and pharmacological inhibition of necroptosis and apoptosis, we show that *ex vivo,* kanamycin and cisplatin ototoxicity is mediated solely by caspase-mediated apoptosis. In mature mice *in vivo*, however, both RIPK-mediated necroptosis and caspase-8-mediated apoptosis contributed to cisplatin and aminoglycoside ototoxicity. Inhibition of necroptosis and apoptosis using pharmacological compounds is thus a clinically tractable strategy of preventing aminoglycoside and cisplatin ototoxicity.

## Methods

### Animal care and handling

The protocol for care and use of animals was approved by the University of Virginia Animal Care and Use Committee. The University of Virginia is accredited by the American Association for the Accreditation of Laboratory Animal Care. C57BL/6 (Bl6) mice used in this study were ordered from Jackson Laboratory (ME, USA). Bl6 mice served as wild-type controls, to match the strain background of the genetically modified mice also used in this study. Mice with null mutations in Ripk3 and Caspase-8 (Salmena et al., 2003; Newton et al., 2004), all on the C57BL/6 background (tested) were bred to produce Ripk3 homozygous/Caspase-8 homozygous (*Ripk3/Casp-8 DOKO*) and Ripk3 homozygous and Caspase-8 heterozygous (*Ripk3 homo/Casp-8 het*) mice, and studied alongside the wild type groups. *Ripk3 KO* mice were provided by Genentech. At the time of treatment, the mice ranged in age from 8-10 weeks. Neonatal mouse pups [postnatal day 3 (P3)–P4] were killed by rapid decapitation, and mature mice were killed by CO_2_ asphyxiation followed by cervical dislocation.

### Organotypic explant cultures

Mouse cochleae and utricles were dissected in Hank’s balanced salt solution (HBSS, Invitrogen, MA) containing 25 mM HEPES, pH 7.5. The organ of Corti was separated from the spiral lamina and the spiral ligament using fine forceps and attached to the bottom of sterile 35 mm Petri dishes (BD Falcon, NY), with the hair bundle side facing up. The dissection medium was then replaced by two exchanges with culture medium (complete high-glucose DMEM containing 1% FBS, supplemented with ampicillin and ciprofloxacin). Prior to experimental manipulation, explants were pre-cultured for 24h, to allow acclimatization to the culture conditions (Francis et al. 2013). Cisplatin (TEVA, MD NDC 0703-5748-11, injectable solution, 1 mg/ml) and kanamycin (Sigma, MO) were dissolved in water. Stock solutions were prepared in DMSO for Nec-1s (0.1 mg/ml, aka RIP1 inhibitor II or 7-Cl-O-Nec-1, MilliporeSigma, MA) and zVAD-FMK (10 mM, Selleckchem, TX). All control cultures were supplemented with the equal amount of the vehicle (DMSO). The organs of Corti were cultured as a whole. Number of experiments (*n*) for quantification of hair cell numbers, activated caspase-3 positive cells and AHA uptake indicates number of organs.

### Immunocytochemistry

Tissues were fixed for 25 min in 3% formaldehyde, washed three times for 5 min each in PBS, and incubated in blocking buffer (PBS containing 1% bovine serum albumin, 3% normal donkey serum, and 0.2% saponin) for 1 h. Organs were then incubated with primary antibody overnight at room temperature in blocking buffer. Organs were washed three times for 5 min each with PBS and incubated with secondary antibodies (fluorophore-conjugated IgGs at 1:100; Invitrogen) and 0.25 µM phalloidin-Alexa 488 (Invitrogen) in the blocking solution for 1–3 h. Finally, organs were washed five times in PBS and mounted in Vectashield (Vector Laboratories, CA). Samples were imaged using Zeiss LSM810 confocal microscopes. The following antibodies were used in this study: mouse anti-MYO7A antibody (Developmental Studies Hybridoma Bank, IA, 1:100), rabbit anti cleaved caspase-3 antibody (Asp175; catalog #9661, Cell Signaling, 1:200).

### Hair cell counts from organ of Corti explant experiments

Hair cells were counted based on MYO7A immunoreactivity over a length of 100 µm of the mid-basal turn of the cochlea, ∼4 mm from the apex (roughly corresponding to the 32 kHz region). Cleaved caspase-3 immunoreactivity was counted in the same area. At least 6 organs of Corti were analyzed for each experimental condition. Exact numbers of organs (*n*) are indicated in the legends.

### Hair cell counts from *in vivo* experiments

Mice were killed after the final ABRs. Cochleae were dissected, openings were created at the base and apex of the cochlea and fixed in 4% paraformaldehyde (PFA) (RT-15720, Electron Microscopy Science, PA) for 24h. After decalcification for 7 days in EDTA solution, all turns of the cochlear sensory epithelium were dissected. Hair cell counting was performed for the organ of Corti using Myo7a immunoreactivity as a marker of hair cell presence. After confocal microscopy, images were analyzed using ImageJ. Hair cells were counted from the apical (0.5-1 mm from apex tip, corresponding to the 6-8 kHz region), mid (1.9 – 3.3 mm from the apex tip, corresponding to 12-24 kHz region) and basal turns (4.7-5.5 mm from the apex tip, corresponding to 48-64 kHz region) of the cochlea. Hair cell numbers were converted and plotted as “% of control”, by normalizing to the hair cell numbers in the corresponding area in vehicle-treated control mice. Organ of Corti from at least 5 mice were analyzed for each experimental condition. Exact numbers of organs (*n*) are indicated in the legends.

### Statistics

For statistical analysis, GraphPad Prism (La Jolla, CA) was used. One-way analysis of variance (ANOVA) was used to determine statistically significant differences between the means of the experimental groups. For pair-wise comparisons, a Tukey post-hoc analysis was performed. *P-*values smaller than 0.05 were considered statistically significant. All *n* in statistical analyses refer to number animals (for *in vivo* experiments) and organs of Corti (for *ex vivo* experiments). All error bars indicate standard deviation (SD).

### Mouse model for cisplatin-mediated hearing loss

To elicit cisplatin-induced hearing loss in the mouse, we followed a protocol established by Li et al. (Li et al., 2011). Mice were injected with 200 mg/kg body weight with the loop diuretic furosemide. 1 h later, cisplatin was injected intraperitoneally (i.p.) at a concentration of 1 mg/kg body weight, in a total of 1 ml saline. This was repeated daily for a total of 3 days. Control mice were injected with vehicle (saline or DMSO/saline mix) only. In groups treated with Nec-1s, the drug was injected i.p. one hour prior to treatment with cisplatin. Our standard dose of 3 mg/kg Nec-1s was chosen from a priori protocols with dosing schedules ranging from 1.2 mg/kg to 5mg/kg (Tristão et al., 2016; Wang et al., 2017).

### Mouse model for kanamycin-mediated hearing loss

To elicit kanamycin-induced hearing loss in the mouse, we followed a furosemide/kanamycin combination protocol (Taylor et al., 2008). Mice were injected with one i.p. injection of 600 mg/kg of Kanamycin followed one hour later by an i.p. dose of 400 mg/kg of furosemide. In treatment groups treated with Nec-1s (3 mg/kg or 0.3 mg/kg), the drug was injected in an i.p. fashion one hour before treatment with kanamycin.

### Auditory Brainstem Response (ABR) Testing

ABR testing was performed on each mouse before initiation of drug treatment, 48 hours post-kanamycin treatment, 10 days post-cisplatin treatment, and 2 weeks following initial posttreatment ABR. Anesthesia with a single intraperitoneal injection of Ketamine at 100 mg/kg was performed prior to testing. The mouse was then placed on a Deltaphase isothermal warming pad (Braintree Scientific) and subdermal needle electrodes (FE-7, Grass Technologies) were inserted as follows: (1) reference electrode was inserted in the mastoid area of the right ear, (2) the active electrode was inserted at the vertex of the scalp, (3) the ground electrode was inserted in the left gluteus. All ABR testing took place in a sound-attenuating box (Med Associates, FL) with digital stimuli (Intelligent Hearing Systems, FL) were delivered to the right ear by a high frequency speaker (model: AS-TH400A) through a plastic tube placed in the ear canal. Stimulus was delivered at 32, 22.4, 16, 11.3, and 8kHz in 1024 sweeps. After testing revealed observable response waveforms, the stimulus was decreased by 10 dB until responses were no longer observed. Repeat testing at 5 dB increments above and below the sound level at which waveforms disappeared confirmed the threshold.

## Results

### Pharmacological inhibition of RIPK1 and downstream necroptosis does not protect from kanamycin- or cisplatin-induced hair cell death ***ex vivo***

The relative contribution of apoptosis and necrosis to drug-induced hair cell death was reported to differ between *ex vivo* and *in vivo* experimental contexts (Forge and Schacht, 2000; Jiang et al., 2006; Rybak et al., 2006). For a comprehensive appreciation of the role of necroptosis and apoptosis in aminoglycoside and cisplatin ototoxicity, we therefore performed both *ex vivo* and *in vivo* experiments. Necrostatin-1s (Nec-1s) is a selective inhibitor of receptor-interacting protein kinase RIPK1, which in conjunction with RIPK3, are the principal mediators of necroptosis (Weinlich et al., 2017). We first conducted a dose-response study of the effect of Nec-1s on kanamycin and cisplatin toxicity in early postnatal organ of Corti explant cultures (*ex vivo)*. Organ of Corti explants were pre-cultured overnight in growth medium, and cultured for another 24 h with 0, 0.2, 1, 3, 10 µM Necrostatin-1s (Nec-1s), in the absence or presence of 100 µM cisplatin or 1000 µM kanamycin. After culture, organs were fixed and processed for immunohistochemistry to visualize F-actin (phalloidin), MYO7A and cleaved Caspase-3 immunoreactivity. Hair cell numbers and the frequency of cleaved Caspase-3 positive cells of Nec-1s treated cultures were comparable to vehicle-treated controls, suggesting that inhibition of RIPK1-mediated necroptosis had no effect on hair cells (**Figure 1A**), and failed to provide any protection from cisplatin and kanamycin induced hair cell death *ex vivo* (**Figure 1B, C**). We next examined the combined effect of pan-caspase inhibition (by zVAD-FMK) and RIPK1 inhibition (by Nec-1s). The ototoxins (cisplatin or kanamycin) and the inhibitors (zVAD-FMK and Nec-1s, individually or in combination) were added simultaneously to the explant cultures. In contrast to Nec-1s, the pan-caspase inhibitor zVAD-FMK at 100 µM concentration was highly effective in preventing both Caspase-3 cleavage and hair cell death in explants exposed to kanamycin or cisplatin (**Figure 1D, E, F**). The completeness of zVAD-FMK-mediated protection against cisplatin- or kanamycin-induced hair cell death, and the lack of protection by Nec-1s, suggested that *ex vivo,* the predominant form of cisplatin and kanamycin-induced hair cell death is mediated by caspase-dependent apoptosis, without contribution from necroptosis.

**Figure 1:**
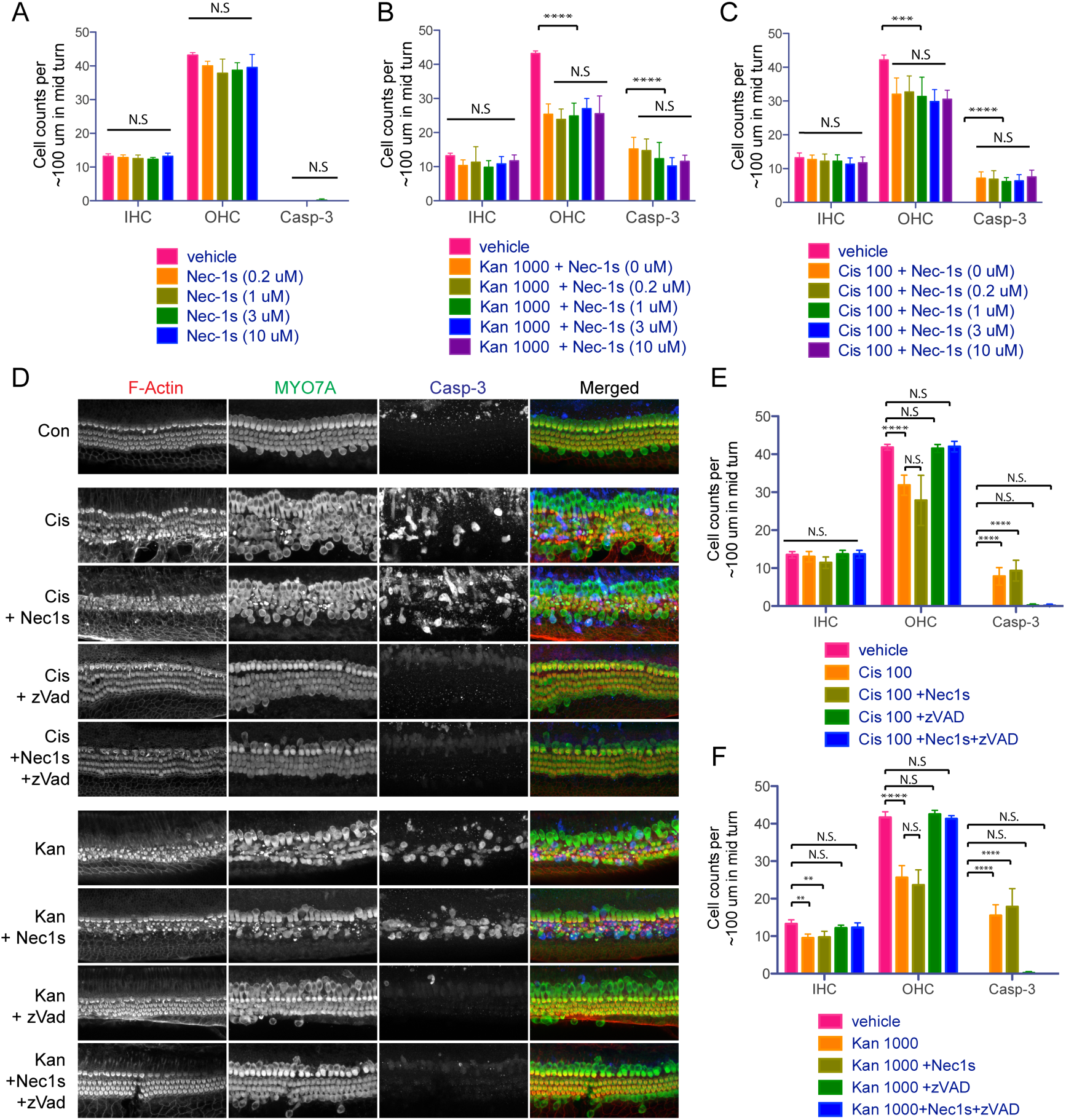
Necrostatin-1s does not protect from kanamycin- or cisplatin-induced hair cell death in early postnatal organ of Corti explant cultures. Dose-response of Nec-1s alone (A), Nec-1s + Kanamycin (Kan) (B) and Nec-1s + Cisplatin (Cis) (C). Counts of inner hair cells (IHC), outer hair cells (OHC) and cleaved caspase-3 immuno-reactive cells in organ of Corti explants exposed to 0.2, 1, 3 or 10 µM Nec-1s, in the absence or presence of 1000 µM kanamycin or 100 uM cisplatin. (D). Representative images of 100 µm long sections in the mid-basal region of cochlear explants under various experimental conditions: Control (con), Cis (cisplatin treated), Cis + Nec-1s (cisplatin and Nec-1s treated), Cis + zVAD (cisplatin and zVAD-FMK treated), Cis + Nec-1s + zVAD (cisplatin, Nec-1s and zVAD-FMK treated). IHC, OHC and caspase-3 positive cells are quantified in E and F. Statistical significance: ****: p-value <0.00001, *** p-value < 0.0001, **: P-value <0.001, *: P-value <0.05. Number of organs (*n*) per group: 6

### Pharmacological blockade of RIPK1 protects against kanamycin-induced hair cell death ***in vivo***

Next, we tested the otoprotective potential of Nec-1s *in vivo*. Previous studies had shown that, in contrast to *ex vivo* cultures, both apoptosis and necrosis contribute to ototoxicity *in vivo* (Jiang et al., 2006). We thus hypothesized that *in vivo*, necroptosis might contribute significantly to kanamycin and cisplatin-mediated ototoxicity. We first tested whether Nec-1s treatment protects against kanamycin ototoxicity in the mouse. Auditory brainstem responses (ABRs) were measured 1 day prior to administration of drugs (pre-exposure ABRs). Nec-1s (3 or 0.3 mg/kg body weight) was administered i.p., followed 1 hour later with a mix of kanamycin and loop diuretics (**Figure 2A)**. Post-exposure ABRs were measured 2 days after drug administration, after which mice were killed and processed for hair cell counts. In contrast to *ex vivo*, RIPK inhibition by Nec-1s provided significant protection from kanamycin induced hearing loss *in vivo*: Control mice treated with a kanamycin/furosemide combination developed hearing loss across all frequencies, with ABR thresholds ranging between ∼70 and 80 dB two days after the kanamycin injection (**Figure 2B, E**). The broad range of affected frequencies is in agreement with previous reports using the combined kanamycin/loop diuretic hair cell damage protocol (Hirose and Sato, 2011). The panels in **Figure 2B-D** indicate absolute ABR thresholds, while the summary plot **Figure 2E** shows the relative ABR threshold difference (shift) between pre- and post-ABRs. In mice that were additionally treated with 3 mg/kg Nec-1s, the thresholds increased only to ∼40-50 dB across tested frequencies, representing an improvement of threshold shift by an average of ∼30 dB in this group (**Figure 2C, E**). When Nec-1s was administered at a 10-fold diluted concentration, the protective effect was lost, demonstrating a dose-dependent protection of Nec-1s in *vivo* (**Figure 2D, E**). To test whether the protective effect of Nec-1s was maintained after withdrawal of the drug, we determined ABR thresholds 14 days (D14) after Nec-1s administration. The improvement in hearing threshold in the Nec-1s treated group was maintained at day 14 (**Figure 2F**), suggesting the beneficial effect of Nec-1s for hearing function persisted after its withdrawal. It should be noted that the ototoxin kanamycin was also absent during this time period. We conclude that *in vivo,* necroptosis contributes significantly to kanamycin-induced hearing loss.

**Figure 2:**
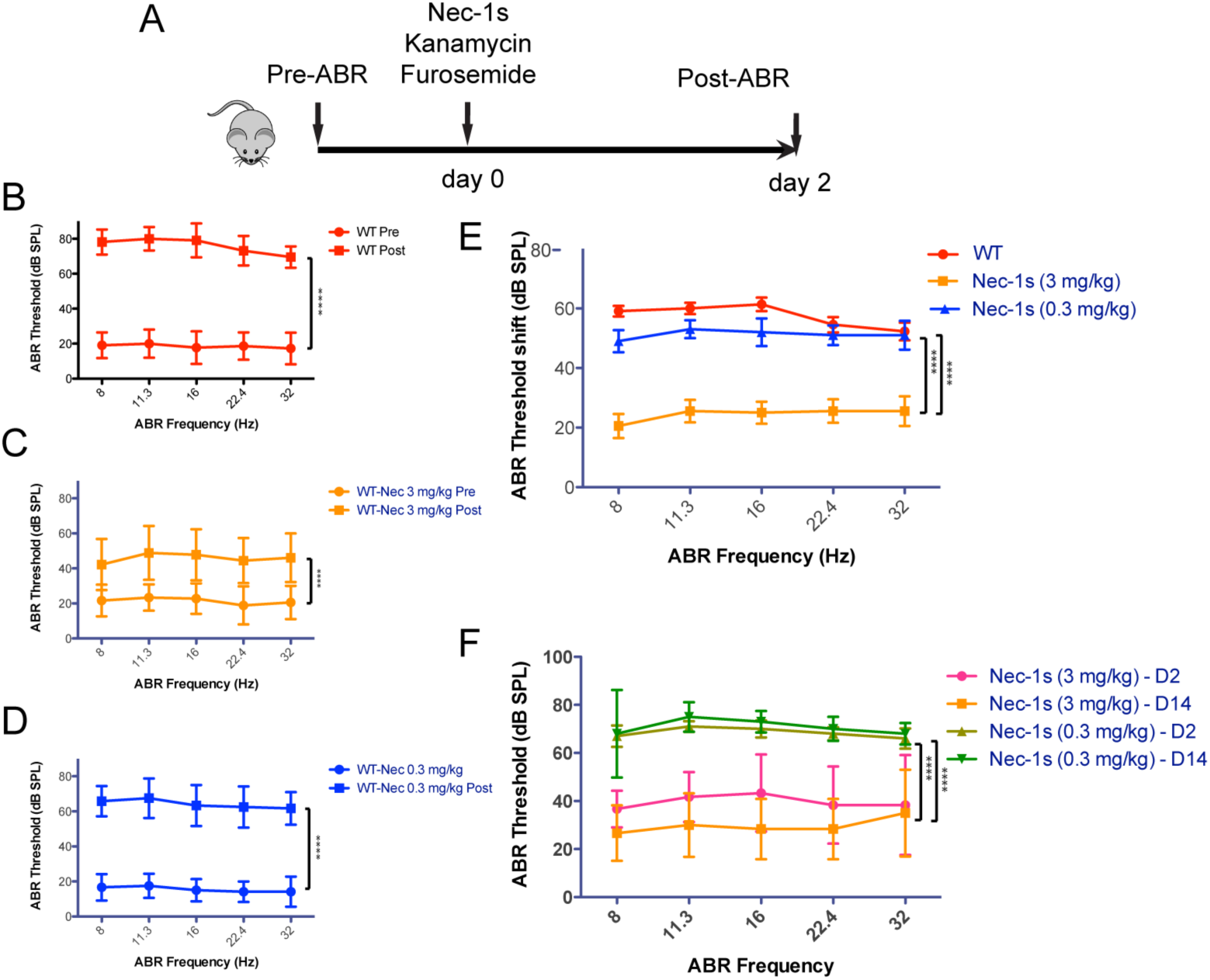
Nec-1s protects from kanamycin induced hair cell death ***in vivo***. Schematic illustration of the kanamycin/furosemide combination ototoxicity paradigm (A). ABR thresholds measured 1-day prior (pre) and 2 days after (post) application of kanamycin/furosemide (according to A) in WT mice injected with vehicle (B), 3 mg/kg body weight Nec-1s (C), and 0.3 mg/kg body weight Nec-1s (D). The shifts (difference between pre and post-exposure ABRs) are summarized for all three experimental treatments in (E). The colors correspond to the respective colors in the individual plots. (F) Follow-up study of ABR thresholds 14 days after exposure. Error bars indicated Standard deviation. Statistical significance: ****: p-value <0.00001, *** p-value < 0.0001. Number of animals (*n*) per group: WT vehicle-treated (*n*=11), WT Nec-1s (3 mg/kg)-treated (*n*=9), WT Nec-1s (0.3 mg/kg)-treated (*n*=5)

### Pharmacological blockade of RIPK1 protects against cisplatin-induced hair cell death ***in vivo***

We next tested whether Nec-1s also alleviates cisplatin-induced ototoxicity *in vivo*. ABRs were tested 1 day prior to administration of drugs (pre-exposure ABRs). Nec-1s (3 or 0.3 mg/kg body weight) was administered i.p., followed 1 hour later with a mix of cisplatin with a loop diuretic (furosemide) (**Figure 3A)**. This was repeated daily for 3 days. Post-exposure ABRs were measured 10 days after the completion of drug administration, after which mice were killed and processed for hair cell counts. Similar to the results for kanamycin ototoxicity, administration of 3 mg/kg Nec-1s provided significant protection from ototoxic hearing loss. ABR thresholds improved an average of ∼30 dB across all tested frequencies in the Nec-1s treated group (**Figure 3B, C, E**). When Nec-1s was administered at a 10-fold diluted concentration, the protective effect was again lost, and hearing thresholds reverted to values similar to WTs exposed to cisplatin (**Figure 3D, E**). The **Figures 3B-D** indicate ABR thresholds, while the summary plot in **Figure 3E** shows the ABR threshold shifts between pre- and post-ABRs. To test whether the protective effect of Nec-1s was maintained after withdrawal of the drug, we performed a separate experiment in which ABR thresholds were determined 24 days (D24) (initial post-exposure ABRs at D10) after Nec-1s administration. The hearing benefit in the Nec-1s treated group was maintained up to 24 days, while the hearing thresholds without Nec-1s treatment even worsened between 10 days and 24 days after cisplatin treatment (**Figure 3F**), suggesting the beneficial effect of nec-1s for hearing function persisted after its withdrawal.

**Figure 3:**
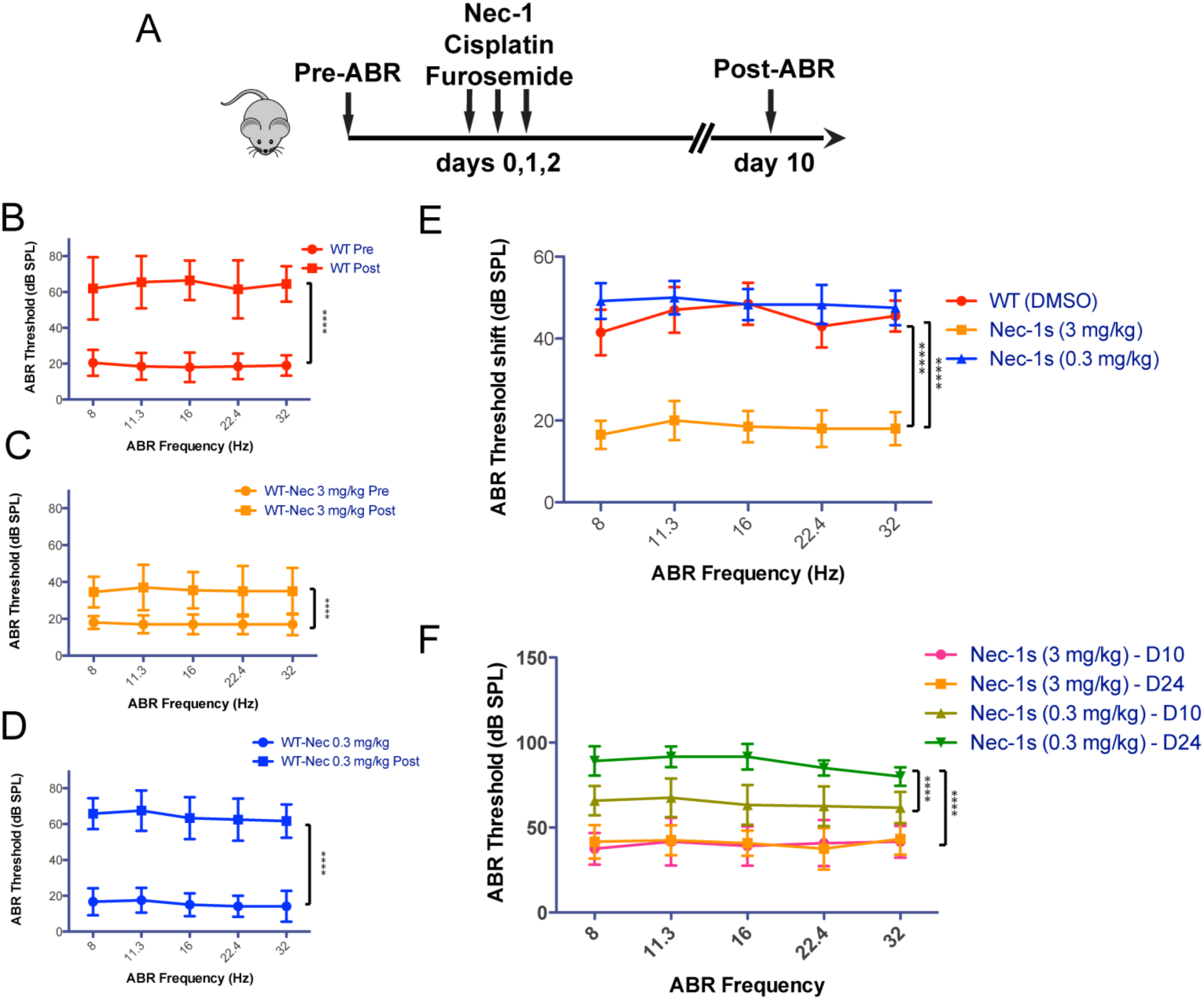
Nec-1s protects from cisplatin induced hair cell death ***in vivo***. Schematic illustration of the cisplatin/furosemide combination ototoxicity paradigm (A). ABR thresholds measured 1 day prior and 10 days after application of cisplatin/furosemide (according to A) in WT mice injected with vehicle (B), 3 mg/kg body weight Nec-1s (C), and 0.3 mg/kg body weight Nec-1s (D). The shifts (difference between pre and post-exposure ABRs) are summarized for all three experimental treatments in (E). The colors correspond to the respective colors in the individual plots. (F) Follow-up study of ABR thresholds 24 days (D24), as compared to the initial post-ABRs 10 days (D10) after exposure. Error bars indicated Standard deviation. Statistical significance: ****: p-value <0.00001, *** p-value < 0.0001. Number of animals (*n*) per group: WT vehicle-treated (*n*=10), WT Nec-1s (3 mg/kg)-treated (*n*=10), WT Nec-1s (0.3 mg/kg)-treated (*n*=6)

Taken together, we conclude that the effect of Nec-1s mediated inhibition of necroptosis differs starkly between *in vivo* and *ex vivo* experiments, and that *in vivo*, pharmacological inhibition of RIPK1-mediated necroptosis alleviates both kanamycin and cisplatin hearing loss.

### *Ripk3/Casp-8 DOKO* **mice exhibit normal ABR thresholds**

Next, we used genetic mouse models to assess the relative contributions of RIPK3 and Caspase-8 in kanamycin and cisplatin ototoxicity *in vivo*. We decided to make use of mice that harbor single mutations in the *Ripk3* gene or combined null mutations in the *Ripk3* and *Caspase-8 (Casp8)* genes. RIPK3, which is recruited by RIPK1 to assemble the necrosome (He et al., 2009; Li et al., 2012) is an essential mediator of necroptosis. Caspase-8 is an “initiator caspase” essential for death-receptor mediated (extrinsic) apoptosis (Chen and Wang, 2002; Boatright and Salvesen, 2003). *Ripk3 KO* (global KO) mice were previously shown to be devoid of any phenotype at basal state (Newton et al., 2004), but protected from various types of stress related responses (Duprez et al., 2011; Ramachandran et al., 2013; Murakami et al., 2014). *Casp8 KO* mice exhibit embryonic lethality, but fortuitously, the embryonic lethality of *Casp8* deletion is mediated by RIPK3, so that mice with a dual deletion of *Casp8* and *Ripk3* are viable and healthy, with the exception of developing lymphadenopathy starting at around 14-16 weeks of age (Kaiser et al., 2011). The present experiments were performed with mice aged 8-10 weeks, to avoid unforeseen interactions with the lymphadenopathy symptoms. To sum up our choice of mouse strains, we used *Ripk3 single KO* mice to study the effect of genetic inhibition of RIPK-mediated necroptosis, and *Ripk3/Casp8 DOKO* to study the combined effect of genetically inhibiting both necroptosis and extrinsic apoptosis on kanamycin and cisplatin mediated ototoxicity. Prior to exposing the mice to the ototoxicity paradigms, we first ascertained that the *Ripk3/Casp8 DOKO* mice have normal hearing function. We performed a longitudinal hearing function study, testing ABRs at 8, 11, 14 and 20 weeks of age in *Ripk3/Casp8 DOKO* and WT controls and found no significant differences between the two groups (**Figure 4 A-D**). We also tested whether the lack of *Caspase-8* might affect the apoptosis of cells in the Kolliker’s organ between postnatal days 7 and 13, thought to contribute to the remodeling of the greater epithelial ridge during postnatal development (Hinojosa, 1977; Knipper et al., 1999; Kamiya et al., 2001). Counting caspase-3 positive cells in the Kolliker’s organ at P8 did not reveal any differences between *Ripk3/Casp8 DOKO* and WT controls (**Fig 4.E, F**). We conclude that *Ripk3/Casp8 DOKO* mice have a likely normal cochlear development and normal hearing function up to 20 weeks of age.

**Figure 4:**
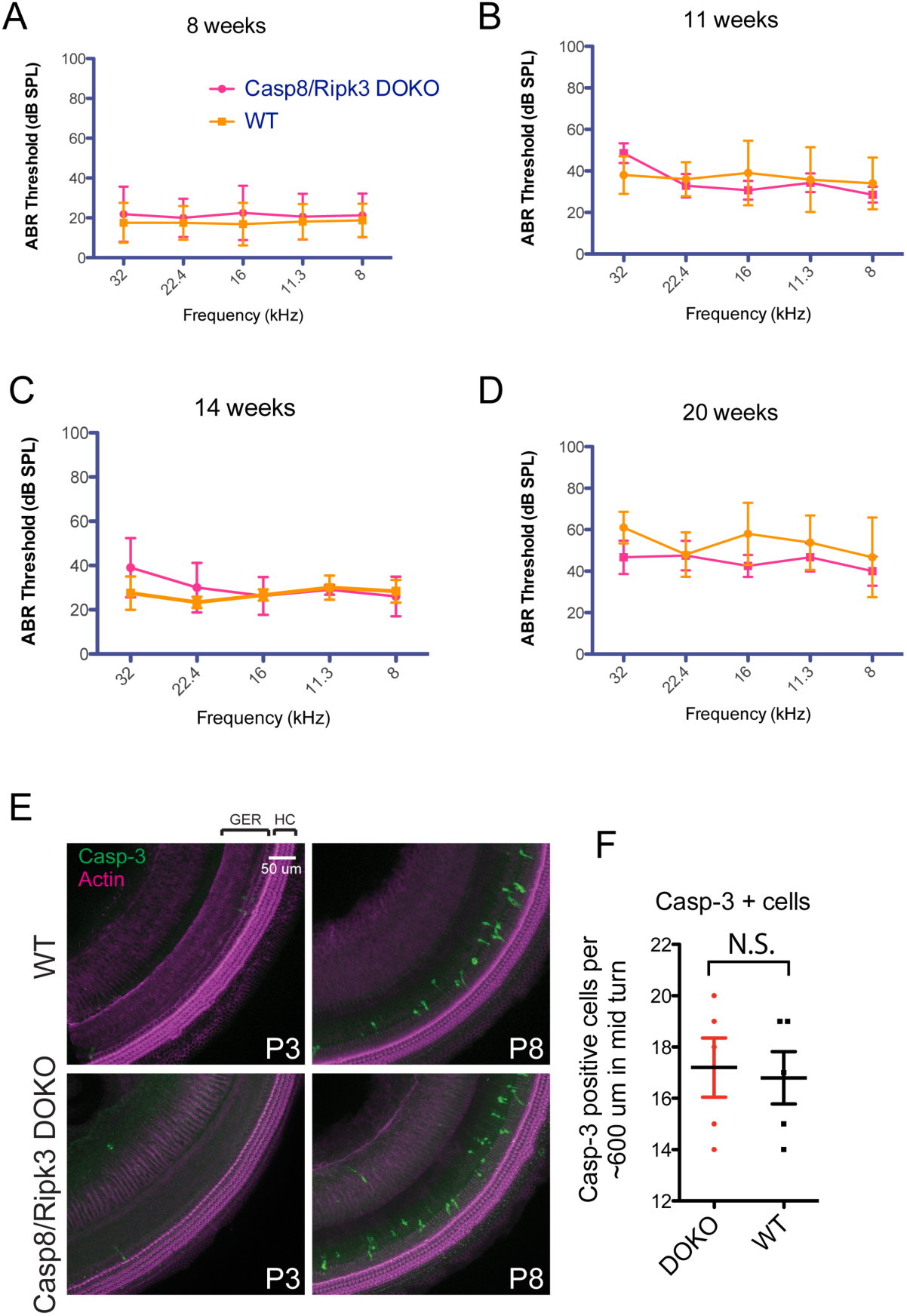
Genetic inhibition of RIPK3-mediated necroptosis and Caspase-8 mediated apoptosis does not affect normal hearing in mice. Comparable auditory thresholds of *Casp8/Ripk3 double-knockout (DOKO)* and WT mice at 8-weeks (A), 11-weeks (B), 14-weeks (C), and 20-weeks (D) of age. Caspase-3 (Casp-3) staining of the mid-turn of the cochlea in *Casp8/Ripk3 DOKO* and WT mice at P3 and P8 of age (E). Cell counts of Casp-3 positive cells in *Casp8/Ripk3 DOKO* and WT mice (F). Number of animals (*n*) per group: WT (8 weeks: *n*=8, 11 weeks: *n*=7, 14 weeks: *n*=6, 20 weeks: *n*=6), *Ripk3/Casp-8 DOKO* (8 weeks: *n*=8, 11 weeks: *n*=6, 14 weeks: *n*=6, 20 weeks: *n*=5)

### Genetic inhibition of RIPK3 and Caspase-8 alleviates kanamycin ototoxicity ***in vivo***

We next tested whether *Ripk3 single KO* mice and *Ripk3/Casp8 DOKO* mice are protected from kanamycin ototoxicity *in vivo*. Mice were exposed to the same kanamycin ototoxicity protocol used for the Nec-1s inhibitor studies (**Figure 5A**). When WT mice were exposed to kanamycin, the hearing thresholds increased by ∼50-60 dB compared to pre-exposure ABR thresholds, affecting all frequencies in a comparable manner (**Fig 5. B, E**). The **Figures 5B-D** indicate absolute ABR thresholds, while the summary plot in **Figure 5E** plots the ABR threshold difference (shift) between pre- and post-ABRs. *Ripk3 KO mice* presented with ABR thresholds that were significantly improved compared to WT mice, with thresholds shifting 2 days after kanamycin exposure by only ∼30 dB in average (**Figure 5C, E**). *Ripk3/Casp8 DOKO* mice were protected most, with ABR thresholds shifting by only ∼15-20 dB after kanamycin exposure (**Figure 5D, E**), indicating that combined deletion of RIPK3-mediated necroptosis and Caspase-8 mediated apoptosis alleviates kanamycin ototoxicity in an additive manner.

**Figure 5:**
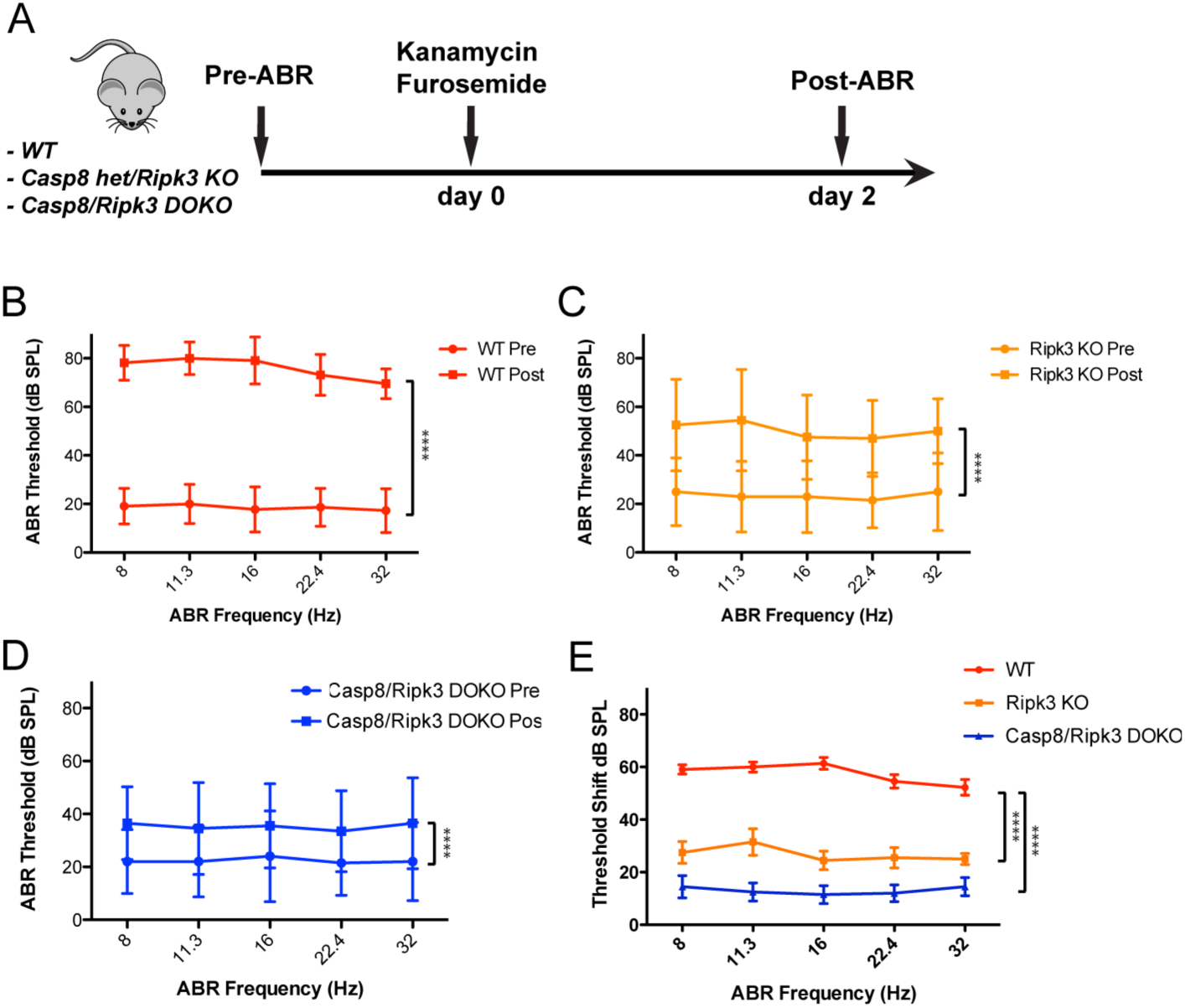
Genetic inhibition of RIPK3-mediated necroptosis and Caspase-8 mediated apoptosis alleviates kanamycin ototoxicity ***in vivo***. Schematic illustration of the kanamycin/furosemide combination ototoxicity paradigm in mutant mice (A). Auditory thresholds increased ∼50-60 dB compared to pre-exposure in WT mice (B). Auditory thresholds increased ∼30 dB compared to pre-exposure in *Ripk3 KO mice* (het for Casp-8) mice (C). Auditory thresholds increased ∼15-20 dB compared to pre-exposure in *Ripk3/Casp8 DOKO* mice (D). Degree of threshold shift at each frequency for each genotype (E). Error bars indicated Standard deviation. Statistical significance: ****: p-value <0.00001. Number of animals (*n*) per group: WT (*n*=11), *Ripk3 KO* (*n*=10), *Ripk3/Casp8 DOKO* (*n*=10)

### Genetic inhibition of RIPK3 and Caspase-8 alleviates cisplatin ototoxicity ***in vivo***

We next tested whether *Ripk3 single KO* mice and *Ripk3/Casp8 DOKO* mice are protected from cisplatin ototoxicity *in vivo*. Mice were exposed to the cisplatin ototoxicity protocol used for the Nec-1s inhibitor studies (**Figure 6A**). WT mice exposed to cisplatin presented with hearing thresholds of ∼60-70 dB, which constitutes a shift of ∼40-50 dB compared to pre-exposure ABRs. All frequencies were affected in a comparable manner (**Fig 6B, E**). *Ripk3 KO mice* presented with hearing thresholds of ∼40-50 dB, which constitutes a shift of only 15-20 dB compared to pre-exposure ABRs, thus were significantly improved compared to WT mice (**Figure 6C, E**). *Ripk3/Casp-8 DOKO* mice experienced the most pronounced protection, with ABR thresholds shifting by only ∼10 dB after cisplatin exposure (**Figure 6D, E**). This indicates that combined deletion of RIPK3-mediated necroptosis and Caspase-8 mediated apoptosis alleviates cisplatin ototoxicity in an additive manner.

**Figure 6:**
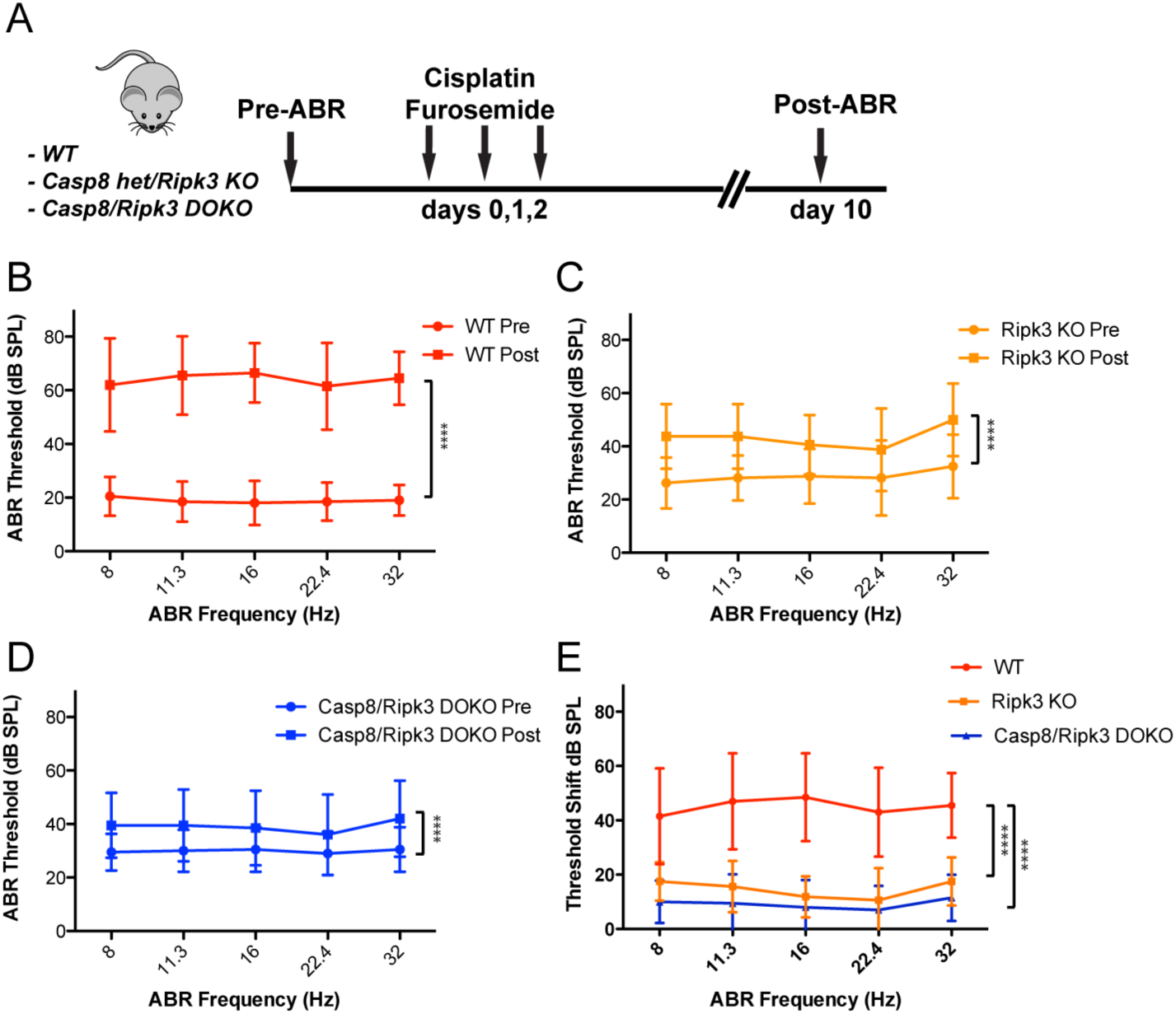
Genetic inhibition of RIPK3-mediated necroptosis and Caspase-8 mediated apoptosis alleviates cisplatin ototoxicity ***in vivo***. Schematic illustration of the cisplatin/furosemide combination ototoxicity paradigm in mutant mice (A). Auditory thresholds increased ∼40-50 dB compared to pre-exposure in WT mice (B). Auditory thresholds increased ∼15-20 dB compared to pre-exposure in *Ripk3 KO mice* (het for Casp-8) mice (C). Auditory thresholds increased ∼10 dB compared to pre-exposure in *Ripk3/Casp8 DOKO* mice (D). Degree of threshold shift at each frequency for each genotype (E). Error bars indicated Standard deviation. Statistical significance: ****: p-value <0.00001. Number of animals (*n*) per group: WT (*n*=10), *Ripk3 KO* (*n*=8), *Ripk3/Casp8 DOKO* (*n*=10)

### The otoprotective effect of inhibiting RIPK3-mediated necroptosis and Caspase-8 mediated apoptosis correlates with improved hair cell survival

If the improved hearing performance in mice treated with Nec-1s or deficient for *Ripk3* and/or *Casp8* is mediated by inhibition of necroptotic and/or apoptotic cell death, it should be reflected in improved hair cell survival. We tested this by counting cochlear hair cells from all experimental groups. Hair cell counts were performed from the apical (0.5-1 mm from apex tip, corresponding to the 6-8 kHz region), mid (1.9–3.3 mm from the apex tip, corresponding to 12-24 kHz region) and basal turns (4.7-5.5 mm from the apex tip, corresponding to 48-64 kHz region) of the cochlea. In WT mice exposed to kanamycin, inner hair cells in the apical and mid regions were mostly unaffected, but basal inner hair cells were significantly reduced in numbers (IHC counts ∼80% lower compared to unexposed control mice) (**Figure 7A, C)**. IHCs in *Ripk3 single KO* and *Ripk3/Casp8 DOKO* mice, and mice treated with Nec-1 maintained most IHCs, with minor (10-20%) losses reported at basal regions compared to untreated controls (**Figure 7A, C**). OHC numbers were much more affected by kanamycin exposure: In WT mice exposed to kanamycin, virtually all OHCs were lost (**Figure 7A, B**). *Ripk3 single KO* mice showed improved hair cell survival, and the OHCs of *Ripk3/Casp8 DOKO* mice were most protected (**Figure 7A, B**). Nec-1s treated mice showed some protection but did not reach the level of *Ripk3 KO* mice (**Figure 7A, B**).

**Figure 7:**
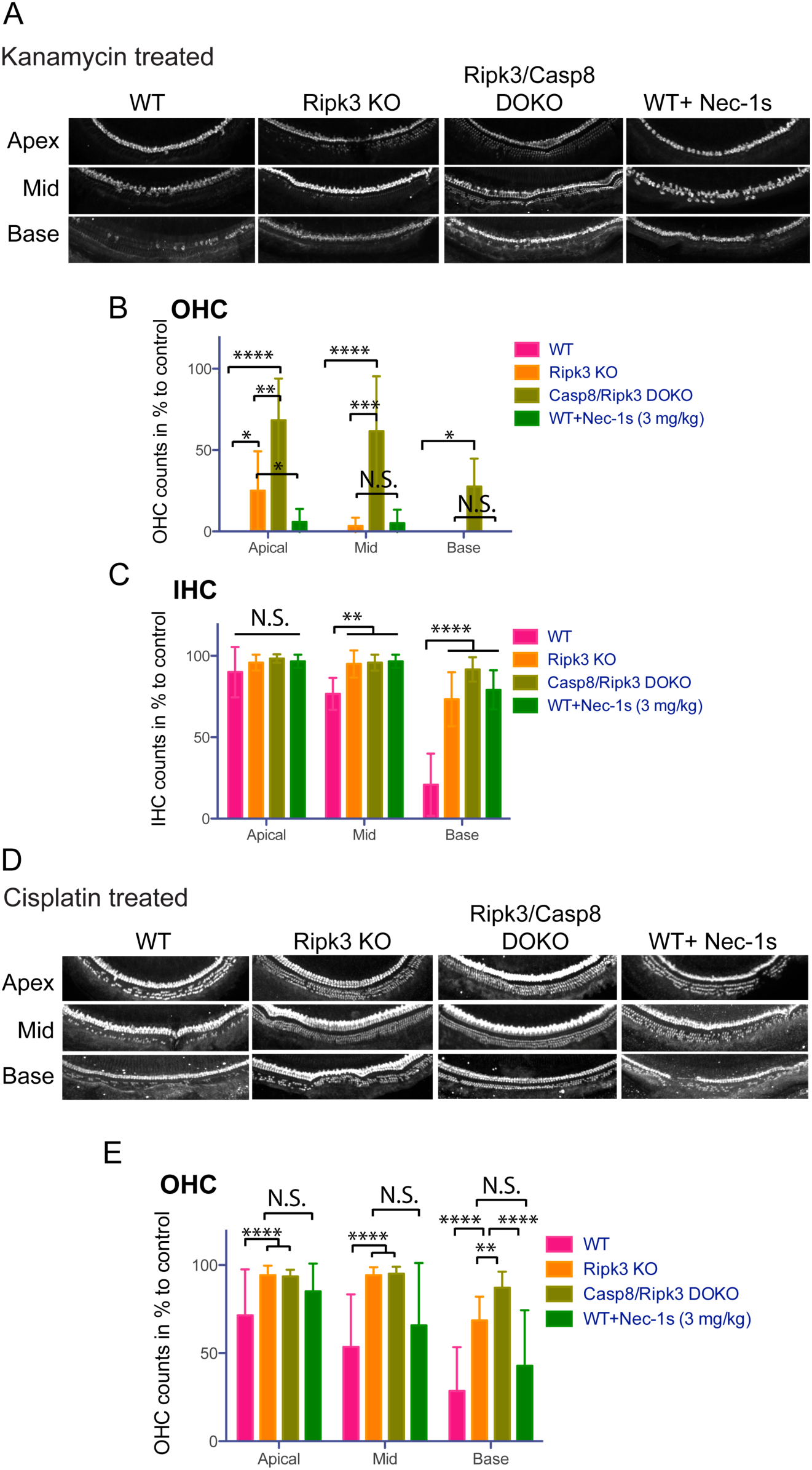
The otoprotective effect of inhibiting RIPK3-mediated necroptosis and Caspase-8 mediated apoptosis correlates with improved hair cell survival. Cell counts of cochleae after treatment with kanamycin stratified by genotype showing a protective effect in *Ripk3 KO, Casp8/Ripk3 DOKO* and WT treated with Nec-1s (A). Outer hair cell counts after treatment with kanamycin stratified by genotype and treatment with Nec-1s (B). Inner hair cell counts after treatment with kanamycin stratified by genotype and treatment with Nec-1s (C). Cell counts of cochleae after treatment with cisplatin stratified by genotype showing a protective effect in *Ripk3 KO, Casp8/Ripk3 DOKO* and WT treated with Nec-1s (D). Outer hair cell counts after treatment with cisplatin stratified by genotype and treatment with Nec-1s (E). Error bars indicated Standard deviation. Statistical significance: ****: p-value <0.00001, *** p-value < 0.0001, **: P-value <0.001, *: P-value <0.05. Number of animals (*n*) per group, only 1 ear per group was analyzed: WT, cisplatin treated (*n*=7), *Ripk3 KO,* cisplatin treated (*n*=7), *Ripk3/Casp8 DOKO,* cisplatin treated (*n*=7), WT, cisplatin-treated + Nec-1s (3 mg/kg)-treated (*n*=7), WT, kanamycin treated (*n*=6), *Ripk3 KO,* kanamycin treated (*n*=7), *Ripk3/Casp8 DOKO,* kanamycin treated (*n*=6), WT, kanamycin-treated + Nec-1s (3 mg/kg)-treated (*n*=6),

In the cisplatin treated group, inner hair cells were not affected at all, in all mouse genotypes (no quantification shown), while OHCs were affected in a genotype-dependent manner (**Figure 7D, E**). In WT mice exposed to cisplatin, OHC numbers were reduced by ∼30, 50 and 70 % (apex, mid, base) compared to untreated controls (**Figure 7D, E**). *Ripk3 single KO* mice showed improved hair cell survival, retaining most hair cells at apical and mid turns, but loosing ∼30% at the base (**Figure 7D, E**). OHC counts in *Ripk3/Casp8 DOKO* mice were similar to the single KO at the apex and mid, but were better at the base **(Figure 7D, E**). Nec-1s treated mice showed some protection compared to WT mice but did not reach the level of *Ripk3 KO* mice (**Figure 7D, E**). In summary, we conclude that the otoprotective effect of inhibiting RIPK3-mediated necroptosis and Caspase-8 mediated apoptosis correlates with improved hair cell survival.

## Discussion

Necroptosis is executed by RIPK1 and RIPK3 when apoptosis-mediating caspases are inhibited (Yuan et al., 2016). Due to this compensatory interplay between the cell death pathways, necroptosis and apoptosis are best studied in mutual context. We achieved this by studying the ototoxic response of mouse models in which RIPK3-mediated necroptosis, and/or Caspase-8 mediated apoptosis is disrupted. In addition, we tested the *ex vivo* and *in vivo* efficiency of the small-molecule inhibitor of RIPK1 Nec-1s.

Historically, both *ex vivo* and *in vivo* studies have provided important insights into the mechanisms underlying hair cell death pathways (Wu et al., 2001; Wang et al., 2004; Rybak, 2007; Rybak et al., 2007; Guthrie, 2008; Warchol, 2010; Ding et al., 2011; Schacht et al., 2012; Francis et al., 2013; Nicholas et al., 2017). The conclusions, however, differ significantly between the *ex vivo* and *in vivo* context, and thus have created confusion in the ototoxicity field. Inconsistencies possibly arise from differences in age (usually P3-4 *ex vivo*, and adult mice for *in vivo* experiments), the denervation of hair cells and absence of endolymph and strial environment in the *ex vivo* context, to name just the most salient differences. In general, apoptotic cell death was more frequently implicated in *ex vivo* studies, while a mix of apoptotic, necrotic and caspase-independent cell death was implicated in *in vivo* studies (Forge and Schacht, 2000; Jiang et al., 2006; Rybak et al., 2006). This is in good agreement with our present study: in our *ex vivo* experiments, two lines of evidence suggest that necroptosis is not involved in hair cell death: First, addition of the necroptosis inhibitor Nec-1s had no effect on hair cell survival, neither in the unstressed nor kanamycin- and cisplatin-exposed explants. Furthermore, the pan-caspase inhibitor zVAD-FMK provided virtually complete protection against both kanamycin- and cisplatin-induced hair cell deaths. If hair cells *ex vivo* were capable of necroptosis, a subset would have found a way to undergo necroptotic cell death despite inhibition of caspases by zVAD-FMK, since inhibition of caspases triggers necroptosis (Zheng et al., 2014; Yuan et al., 2016). It is thus likely that in the *ex vivo* context, most, if not all hair cell death can be attributed to caspase-mediated apoptosis. This is in agreement with a multitude of previous studies (Forge and Schacht, 2000; Cheng et al., 2005; Jiang et al., 2006; Rybak et al., 2006; Rybak and Ramkumar, 2007; Huth et al., 2011; Schacht et al., 2012; Sheth et al., 2017). *In vivo*, however, our study found that both necroptosis and apoptosis contribute to kanamycin and cisplatin ototoxicities. Previous studies have provided evidence for an involvement of apoptosis *in vivo*: In chick, Matsui et al. showed that infusion of zVAD-FMK into the inner ear protected from streptomycin-induced hair cell death (Matsui et al. 2003). Similarly, infusion of z-DEVD-FMK provided protection from cisplatin-induced ototoxicity in Guinea pigs *in vivo* (Wang et al., 2004). In contrast, Jiang et al. reported that in mice exposed to kanamycin, hair cell nuclei exhibited both apoptotic- and necrotic-like appearances, but lacked molecular hallmarks of classic apoptotic pathways (caspases, cytochrome C, TUNEL) (Jiang et al., 2006). Our use of *Casp8 KO* mice (on a *Ripk3 KO* background) provides first time genetic evidence that Caspase-8 mediated form of apoptosis, also known as death receptor mediated or extrinsic apoptosis, is involved in cisplatin- and kanamycin-mediated ototoxicity. The involvement of the other arm of apoptosis, mitochondrial (intrinsic) apoptosis, was not tested in this study, but previous studies using Caspase-9 inhibitors indicate that the intrinsic apoptotic pathways in implicated in various ototoxicities (Wang et al., 2003). To date, with the exception of the present study, only one other study has explored the role of necroptosis in any type of ototoxicity: Zheng et al. reported that necrostatin alleviates noise-induced hearing loss in the mouse (Zheng et al., 2014). Our study extends the significance of necroptosis to drug-induced ototoxicity, using both pharmacological and genetic intervention.

In general, inhibition of necroptosis and/or apoptosis had similar protective effects on kanamycin and cisplatin ototoxicity. A few subtle, but informative differences were noted: As seen in **Figures 6 and 7,** compound deletion of both *Ripk3* and *Caspase-8* provides increased protection from kanamycin ototoxicity compared to single deletion of *Ripk3*, suggesting that RIPK3-dependent necroptosis and Caspase-8 mediated apoptosis operate in an additive manner (**Figure 5E**). In cisplatin ototoxicity, however, single deletion of *Ripk3* provided nearly equal protection compared to *Ripk3/Casp8* compound deletion, suggesting that i) necroptosis plays a relatively bigger role in cisplatin ototoxicity, and ii) that Caspase-8-mediated apoptosis is less relevant for cisplatin ototoxicity. This is in agreement with a previous study showing that infusion of the Caspase-8 specific inhibitor z-IETD-FMK into Guinea pigs failed to provide protection from cisplatin-induced hearing loss (Wang et al., 2004).

In our study, hearing rescue correlates with improved hair cell survival, which is consistent with the hypothesis that hair cells are the mediators of ototoxicity and otoprotection. However, it is also possible that the benefit of necroptosis and apoptosis inhibition is not mediated by hair cells, and that the improvement of hair cell survival is a secondary effect, possibly by protection of other cell types such as strial cells. This is especially relevant considering the fact that the ototoxicity protocols used in this study, utilizing loop diuretics to increase penetration of ototoxins through the blood-labyrinth barrier, is known to cause additional damage to the stria vascularis (Hirose and Sato, 2011). The question of the primary target cell type of cisplatin and aminoglycoside ototoxicity needs to be addressed in cell-type specific studies using conditional KO mice.

In contrast to apoptosis characterized by its traceless removal of dead cells, necroptosis, due to release of intracellular components acting as damage-associated molecular patterns (DAMPs) in the extracellular space, creates an inflammatory response, which is often beneficial for resolving the insult. A hair cell with necroptotic response thus would only make sense if the inner ear was not “immune privileged” as historically assumed but was capable of mounting a significant immunological response. In fact, more recent studies suggest that the resolution of ototoxic insults is aided by immune cells infiltrating into the inner ear (Kaur, 2015; Kaur et al., 2015, 2018; Francis and Cunningham, 2017; Mizushima et al., 2017; Wood and Zuo, 2017; Barald et al., 2018; Liu et al., 2018). Such immune activity, beneficial under normal circumstances, might become counterproductive when ototoxins overwhelm the system, creating a massive, detrimental immune response. With the potential involvement of the immune response in ototoxicity, the interpretation of our study is complicated by another connection to the neuro-immune angle: RIPK3, previously exclusively implicated in necroptosis, was also shown to be important for inflammatory signaling (Daniels et al., 2017). *Ripk3 KO* mice thus might have aberrant inflammatory signaling that might modify, in a positive or negative manner, the otoprotection provided by the inhibition of necroptotic hair cell death. A potential role of RIPK3 in ototoxicity independent from its role in necroptotic cell death could be explored using mice that are deficient in other mediators of necroptosis, such mixed lineage kinase domain-like protein MLKL, which is a downstream effector of RIPK3 (Cho et al., 2009; Wang et al., 2014).

It should be noted that the protection provided by the genetic inhibition of RIPK3-mediated necroptosis, and Caspase-8 mediated apoptosis is incomplete, suggesting that other pathways are involved. For instance, the mitochondrial form of apoptosis is still active in the *Ripk3/Casp8 DOKO*, possibly explaining part of the incomplete protection.

The present study provides insight into the basic mechanisms underlying drug-induced hearing loss, and importantly, opens up avenues for pharmacological intervention towards clinical therapy. The potential clinical uses of the small molecule inhibitor RIPK1 Nec-1s, which can pass the blood-brain barrier, are already being investigated for neurodegenerative diseases (Zhang et al., 2017), and clinical trials are underway for other necroptosis inhibitors (Weisel et al., 2017). It seems prudent that this class of medication may be translated to clinical studies to assess its otoprotective capabilities.

## Conclusion

The clinical application of cisplatin and aminoglycosides is limited due to ototoxic side effects. Two types of programmed cell death, apoptosis and necroptosis, contribute to aminoglycoside and cisplatin ototoxicity. Direct inhibition of the initiators of apoptosis and necroptosis is possible. This presents new avenues for pharmaceutical intervention to prevent ototoxic hearing loss.

